# The double-edged sword of wildlife biodiversity in the agricultural matrix: a case study of reptiles in vineyards

**DOI:** 10.1101/2025.11.19.689282

**Authors:** Liran Sagi, Akiva Topper, Oded Keynan, Amos Bouskila, Oren Kolodny

## Abstract

The global pervasiveness of agriculture and its profound effects on biodiversity increasingly mandate developing sustainable agricultural methods to support healthy ecosystems within agricultural landscapes. However, increasing the attractiveness of agricultural landscapes for wildlife may negatively impact biodiversity if animals or their reproduction are adversely affected in such habitats. While the effects of agricultural practices were investigated across various crop types and wildlife species, reptiles remain largely overlooked. Additionally, although reptiles in croplands were monitored throughout the farming season, little attention has been directed towards the effects of harvest at the season’s end.

To illustrate the potential impacts of mechanical harvest on wildlife in permanent crops, we present a case-study concerning reptiles in vineyards. We scanned the waste of a medium-sized winery over four nights and found 105 reptiles, 69% of them chameleons. Though chameleons are considered carnivorous, approximately half had consumed grapes, potentially attracted by them. Vineyards, like many crops, offer resources including sugar, water, and shelter, likely attracting many wildlife species. However, mortality during mechanical harvest may be high, potentially rendering such crops an ecological trap. For chameleons the effects may be particularly devastating, as harvest closely precedes the egg-laying season. While our case study focuses on reptiles in vineyards, its implications extend to many species in permanent crops. Ultimately, ecological traps affect biodiversity by attracting animals to habitats in which their fitness is impaired. While preference has been widely discussed in determining trap severity, our case study highlights that considering the trap’s fitness impact is also crucial.

**Highlights:**

- Agricultural landscapes may attract wild animals while increasing their mortality
- Monitoring biodiversity in agriculture must account for the effects of harvest
- Chameleons are likely attracted to vineyards but killed just before reproducing
- Agricultural methods and life history interact to determine ecological trap severity
- Ecological trap severity is affected both by preference and by fitness effects

## 1. Introduction

Since the beginning of the 21^st^ century, agricultural land use has grown by six percent and currently occupies ∼37% of global land area (FAO 2023). Given the global prevalence of agriculture and its profound effects on many ecosystems, much research is directed at devising sustainable methods to combine agricultural productivity with conservation goals. Encouraging the dual use of agricultural landscapes can facilitate the increased presence of natural fauna, benefiting species’ conservation and the ecological system’s health, as well as farmers who may benefit from various ecosystem services. Examples can be found in practices such as planting cover crops to enhance habitat connectivity (e.g., Schipanski *et al*. 2014); agroforestry systems that combine trees and crops to preserve natural habitats alongside agricultural land use (reviewed by Jose 2009); reducing the use of chemical pesticides by encouraging/introducing natural enemies of common pests (reviewed by Zhou *et al*. 2024) etc. The existence of many species of wildlife in agricultural landscapes is generally considered a positive phenomenon in the context of conservation, and even an achievement actively pursued (e.g., Carpio *et al*. 2017). Furthermore, agricultural landscapes are often considered by policymakers to serve as ‘ecological corridors’, bridging between natural patches and allowing gene flow between separate populations (e.g., Rothschild 2011).

However, it is important to address the various ways in which agriculture, including supposedly sustainable agricultural practices, may act as a ‘double-edged sword’, for example, by reducing the fitness of animals attracted to the agricultural landscape. Among animals’ many life-history ‘choices’, habitat selection is often pivotal for an individual’s fitness. To maximize their fitness, it is crucial that animals correctly select suitable sites in which to forage, reproduce, find shelter, etc. However, evaluating the quality of a given habitat may not always be straightforward. Hence, many species have evolved to use various cues that have proven to be good indicators of a habitat’s quality throughout the evolutionary history of the animal’s lineage. For example, certain cues may be associated with an abundance of food or with the presence of suitable sites for reproduction, etc.

With the global expansion of agricultural land use increasingly affecting habitats and species worldwide, a notable risk is that cues used by animals to evaluate habitat quality may no longer be informative or may even be negatively correlated with the habitat’s true quality. This may lead individuals to select habitats resulting in lower fitness than they could otherwise have obtained, in a phenomenon termed an ‘ecological trap’. Ecological traps have been divided into two types: (I) In an ‘equal preference trap’ (Hawlena *et al*. 2010), animals fail to discern the low quality of a habitat and are therefore equally likely to select it relative to other available habitats; (II) In a ‘severe trap’, cues incongruent with the low quality of a habitat result in individuals actively *preferring* low-quality habitats relative to other available possibilities (Hale & Swearer 2016; Robertson & Hutto 2006). Severe traps are considered particularly detrimental, as they are likely to create strong source-sink dynamics that can cause adjacent populations inhabiting high-quality habitats to collapse.

On the one hand, agricultural landscapes may provide certain species with a predictable supply of food and water (including when resources in the natural environment may be scarce); vast areas in which intraspecific and interspecific competition may be reduced; a refuge from certain types of predators, etc. (García-Vega *et al*. 2024). On the other hand, agricultural landscapes may present various threats such as increased predation risk upon crossing between agricultural and natural habitats or within the agricultural habitat itself (e.g., Hansen *et al*. 2019); use of pesticides and other pest control methods (reviewed by Wan *et al*. 2025); and eventual crop harvest. For example, wheat fields have been demonstrated to attract *Heremites vittata* (formerly known as *Trachylepis vittata*), skinks from adjacent remnant patches, but upon harvest of the field, the entire population of *H. vitatta* disappears from the field, presumably killed in the process. Skinks moving from natural patches into wheat fields, or already inhabiting the fields, were found to be in superior body condition relative to skinks in natural patches. Hence, it was demonstrated that a preference for the wheat fields exists, rather than subordinate individuals being competitively excluded into them. Furthermore, juveniles were found only in the natural patches, indicating that reproduction occurs in the natural habitats but not in the wheat fields (Rotem *et al*. 2013).

Importantly, agricultural landscapes that are beneficial to certain species may act as an ecological trap for others; what serves as an ecological corridor between populations of certain species might act as a sink for other species, potentially causing the decline not only of the population within the agricultural area but also of populations adjacent to it. Similarly, different cultivation practices may influence how specific crops affect the species inhabiting the areas in which they are grown. Hence, it is crucial to consider the effects of agriculture on a species-specific, crop-specific, and cultivation practice-specific basis. Between the years 2000-2021, land use by permanent crops (used for long time periods after planting) increased by 40%, in comparison to temporary crops (replanted following each harvest), which increased by 12% during the same time period (FAO 2023). Hence, it is particularly important to consider whether permanent crops, in addition to temporary crops (e.g., Gameiro *et al*. 2024; Rotem *et al*. 2013), may constitute ecological traps.

Vineyards, particularly those serving the winemaking industry, are a popular permanent crop throughout many parts of the world, particularly in Europe. For example, in the EU, vineyards occupy more than three million hectares of land, representing 45% of all vineyards serving the winemaking industry globally. Of these, 83.3% of vineyards occupy less than 1 ha of land (eurostat 2022). Receiving irrigation throughout the growth season (in EU, May-September), vineyards provide available water and shelter when the surrounding environment may offer very little. A common method of harvesting grapes for wine uses machines to shake the vines and dislodge grapes, which are then collected onto a conveyor belt. Following harvest, which typically occurs at night, the grapes, still attached to their stems, are transported to a winery. In the winery, the grapes pass through a machine that destems them. At this stage, the grapes are transferred to the crushing machine while stems and other waste products are discarded. This mechanical harvesting method is faster and more cost-effective than traditional methods, making it attractive for large vineyards (Kurtural & Fidelibus 2021). However, many animals may also be collected during this process. The structural composition offered by vineyards seems to be well suited to arboreal reptiles while the conditions created by the vineyard may attract numerous invertebrates that can serve as potential prey. In addition, some reptiles may also consume fruit and other plant matter found in the vineyards. However, towards the end of the summer, precisely when vineyards are expected to be most attractive compared to the dry surrounding environment, the grapes are harvested, potentially resulting in extreme mortality rates for animals unable to escape.

One avenue for investigating the effects of mechanical harvest on arboreal reptiles in vineyards is to examine the waste discarded following the destemming process. In this manuscript, we present and discuss the findings collected from the waste tank of a medium-sized winery in central Israel.

## 2. Material and methods

### 2.1. Winery survey

To investigate whether reptiles and other animals are accidentally collected during the harvest, we examined the content of the organic waste container in a medium-sized winery in central Israel over four nights in September 2021 and 2023. The aforementioned winery receives grapes from all over the country, ranging from the Negev Desert (Mitzpe Ramon; 30.61°, 34.80°) to the Upper Galilee (33.03°, 35.33°). Grapes processed at this winery are mainly harvested mechanically, typically at night. The organic waste container collects waste following the destemming process, including all non-grape material collected during the harvest. Over the four survey nights, two or three observers stood inside the waste container with flashlights, scanning the waste continuously as it entered the container from the conveyor belt above and collecting any vertebrates found. The winery received and processed grapes from various origins, hence the animals collected were categorized according to the origin of the truck that brought the grape batch in which they were found. At the end of each survey night, we obtained information from the winery, including the size of the vineyards. Dead animals were placed in plastic bags labeled with their location of origin and frozen at -80°C. Live animals were released the following day under the instruction of the Israel Nature and Parks Authority in a designated location.

### 2.2. Density estimations

During the survey nights, trucks originating at the same vineyard did not always arrive sequentially, and in certain cases, we were unable to survey all the batches from a single vineyard. To assess the individual-to-area density of chameleons and agamids collected, we assumed that the yield (mass-to-area ratio) remains constant in the vineyard and used the grape mass to estimate the size of the area harvested. In cases in which vineyard size was unknown, we used the mean mass-to-area ratio (9658 kg/ha) to transform from mass to area. Additionally, in many cases, several grape batches were processed continuously, sometimes rendering it difficult to determine the exact origin of grapes and animals with high certainty. In such cases, individuals were labeled with a broader geographic origin, which was used to assess overall density but not for assessing density at specific locations.

### 2.3. Dissection

Most animals collected during the survey were chameleons, followed by agamids. Hence, we decided to focus on chameleons and dissected them to analyze their stomach content and reproductive status at the time of harvest. Chameleons were thawed prior to dissection, and measurements including mass, tail length, snout-vent length, and head size were recorded for each animal. The sex of each chameleon was determined using secondary sexual characteristics. Each individual was carefully opened, the digestive system was removed, and the number and mass of the eggs were recorded for females. The stomach was opened, and its contents were documented by categorizing food items as either grapes or arthropods. In cases where the stomach did not contain grapes, the intestine was examined for the presence of grape seeds. The digestive system and eggs were placed in separate tubes, labeled for future study, and returned to -80°C.

### 2.4. Analysis

To estimate whether chameleons are drawn to vineyards (i.e., exhibit a preference) we examined several indices of body condition. In chameleons, the number of eggs is correlated with female body size (Díaz-Paniagua et al. 2002) and represents a measure of female condition. We compared the number of eggs found in females collected at the winery to the records from Díaz-Paniagua et al. (2002) which were constructed using WebPlotDigitizer (Rohatgi 2015). The statistical comparison was performed using a permutation test (because in each comparison the values of one group did not distribute normally, p>0.5, Shapiro-Wilk normality test). The effect size was calculated using Cliff’s delta (Cliff 1993).

We also compared the body condition of chameleons collected at the winery to chameleon sizes from a previous study (Bar-Yaacov et al. 2015). Chameleons from the winery were collected during September, while the chameleons measured by Bar-Yaacov et al. (2015) were measured in July. Males and females were compared separately. Although these two datasets represent the same species, there is considerable variation over time in body condition (see appendix 1). It was previously suggested that the mass to snout–vent length (SVL) ratio is the best noninvasive method to assess body condition in lizards (Sion et al., 2021). However, in chameleons, SVL measurements in live individuals exhibit great variability due to behavioral factors such as stress (Pers. Obs,). Therefore, SVL is less reliable as a predictor of body size when comparing live individuals. However, tail length strongly correlates with SVL in chameleons (Sagi *et al*. In review), and they do not autotomize their tails rendering tail length a reliable measurement. Hence, we used the tail-to-mass ratio as a predictor of body condition. The statistical comparison was performed using a permutation test. All comparisons and tests were performed with scripts we wrote in the R environment (R Core Team 2023).

## 3. Results

### 3.1. Winery survey

Over the four nights of the survey, we found 105 reptiles and three individuals of Savigny’s treefrog (*Hyla savignyi*). Among the reptiles, Mediterranean chameleons (*Chamaeleo chamaeleon recticrista*) were the most abundant (72 individuals), followed by 28 Egyptian rock agamas (*Laudakia vulgaris*), two Lebanon lizards (*Phoenicolacerta laevis*), two Mediterranean thin-toed geckos (*Mediodactylus orientalis*), and one dice snake (*Natrix tessellata*). Most of the reptiles were found dead, although 13 chameleons and 3 agamas survived the process.

### 3.2. Density estimations

The density of animals collected at the winery ranged between 0.42 and 6.2 individuals per hectare (ind/ha) for chameleons, and 0 to 7 ind/ha for agamids. Overall density across the respective distribution range was 1.9 ind/ha for chameleons and 0.8 for agamids (see appendix 2). However, it should be noted that there was substantial variation between vineyards. For example, one vineyard in the Judean Mountains contained 6.4 chameleons/ha whereas another vineyard from the same region contained only 1.25 chameleons/ha.

### 3.3. Dissection

We found that consumption of grapes was common in *C. chamaeleon recticrista* individuals originating in vineyards: grapes were found in the stomachs or intestines of 28 individuals. That is, at least 47% (28/60 dead chameleons) had consumed grapes shortly before they were collected. The digestive systems of some additional chameleons contained purple matter that may have been grape residue. However, due to the absence of seeds, we were unable to confirm that this did indeed reflect grape consumption, and therefore these cases were conservatively not included in our count.

Almost all female chameleons contained eggs: we found a total of 404 eggs (range: 3-28) in 24 females. Only one female did not contain eggs, while in another, the eggs were crushed and impossible to count. Another female contained three eggs but was severely damaged prior to collection and was possibly missing some of her clutch. These three chameleons were excluded from comparisons of clutch size. The average clutch size of chameleons found in the winery was larger than previously reported for this species (average in Spain, p = 0.02, permutation test, Cliff’s delta = -0.43, 95% CI: -0.67 to -0.17) from Spain (Díaz-Paniagua et al. 2002).

The body condition of the chameleons was calculated according to the mass-to-tail ratio and we used a permutation to compare our results to the chameleons measured by Bar-Yaacov et al. (2015) in a previous study in Isreal. Body condition in females did not differ between those found in the winery and those surveyed previously in Israel (p = 0.16, Cliff’s delta = -0.17, 95% CI: -0.53 to 0.2). However, the body condition estimates of males were lower in individuals collected at the winery (p = 0.01, Cliff’s delta = 0.52, 95% CI: 0.17 to 0.8).

## 4. Discussion

Alongside the numerous ways in which agriculture may negatively impact the environment, certain agricultural practices may mitigate at least some detrimental effects and even benefit various species of conservation interest. However, it is important to consider the ways in which even ostensibly ‘sustainable’ practices may negatively affect biodiversity. For example, the use of nesting boxes in order to encourage barn owls (*Tyto alba*) as biological agents for pest control in agricultural fields has been demonstrated to have detrimental effects on endangered species of rodents in adjacent natural habitats (ZaitzovelJRaz *et al*. 2020). Similarly, uncultivated remnant patches are generally considered to positively affect biodiversity by enabling various species to persist within the agricultural matrix. However, if animals are unable to reliably assess the quality of natural versus agricultural habitats, agriculture may become an ecological trap, depleting surrounding patches of their natural populations (Rotem *et al*. 2013).

Vineyards, due to their rapid expansion in recent years, have attracted much research attention in the context of nature conservation. A review article from 2020 (Paiola *et al*. 2020) surveyed 218 publications concerning the connections between vineyards and biodiversity conservation. Yet, none of the papers they found concerned reptiles. Furthermore, their review seems to explore the effects of vineyards on biodiversity in and around them, but does not explicitly relate to the effects of harvest. Additional publications (Biaggini & Corti 2021; Kazes *et al*. 2020) specifically examined the abundance of reptiles in vineyards, yet also in these cases, the effects of harvest remained uninvestigated. Biaggini & Corti (2021) reported a stable presence of lizards in vineyards and stressed the importance of investigating the effects of management activities, including mechanical harvest. Policy makers, at least in certain cases, regard vineyards as ecological corridors (e.g., Rothschild 2011), while in other cases strips of remnant vegetation are left within and between vineyards to serve as ecological corridors (Díaz-Forestier *et al*. 2021; Hilty & Merenlender 2004). In general, various recommendations have been made in order to improve the habitat quality for reptiles in permanent crops (Biaggini & Corti 2015; Carpio *et al*. 2017). However, it is crucial to consider not only the presence and abundance of reptiles in vineyards, but also the fitness of reptiles within the vineyards. Attraction of reptiles to mechanically harvested crops may create an ecological trap, decimating the population within the vineyard and potentially endangering populations of reptiles also in adjacent natural patches.

Chameleons constituted almost 70% of the reptiles we collected from the waste pile at the winery (Fig. 1). There may be several reasons for their high representation in our data: first, harvest occurs during autumn, when the surrounding area is mostly dry. Vineyards, in contrast, are irrigated and therefore green and lush and may attract an abundance of potential prey (Chaperon *et al*. 2022). Second, though chameleons are typically regarded as obligate carnivores (but see Keren-Rotem *et al*. 2006), our results demonstrate that, surprisingly, when found in vineyards, grapes are included in their diet (at least 47% of the chameleons we dissected contained grape remnants in their digestive systems). Hence, it is possible that chameleons are actively drawn to vineyards, where grapes may constitute an important part of their diet and provide sugars and fluids in an otherwise hot and dry season. Third, of all the reptiles collected at the winery, only the chameleons and *M. orientalis* are truly arboreal and often spend the nights on vegetation. However, *M. orientalis* hide in crevices and under exfoliating pieces of bark, and are therefore less likely than chameleons to be collected by the harvest machine.

**Figure 1.**
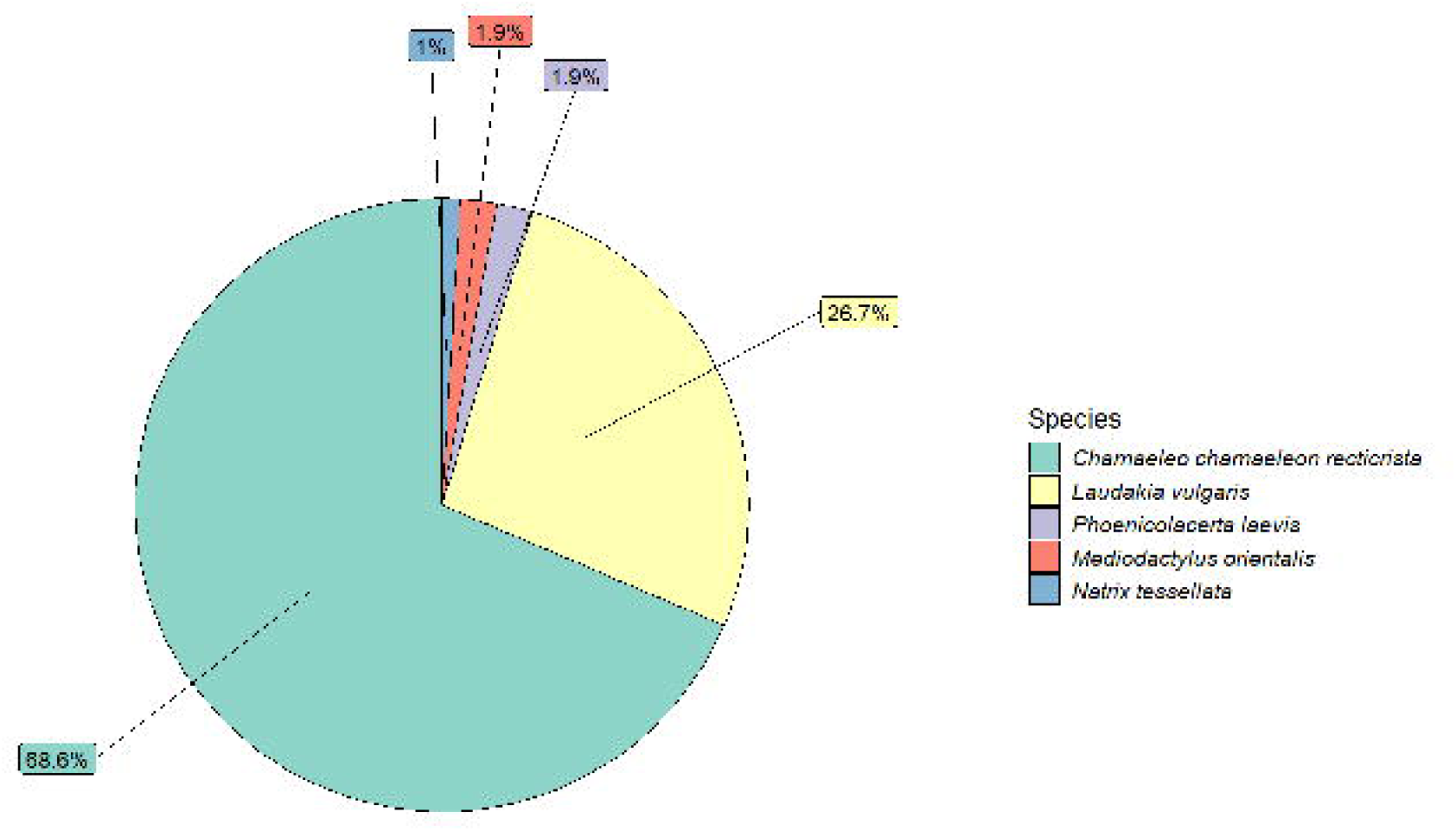
Proportion of reptiles collected from the winery’s waste tank (colored according to taxon).

It has been suggested that three criteria should be fulfilled in order for an ecological trap to be determined (Hale & Swearer 2016; Robertson & Hutto 2006): (I) Individuals prefer the ecological trap habitat (a ‘severe’ trap) or show equal preference for both habitats (an ‘equal preference trap’); (II) Fitness of the individuals, or a reasonable surrogate of fitness, differs between the two habitats; (III) Fitness is lower when animals exploit the ecological trap. While directly determining preference is often impractical, given the two reasons provided above, we consider it likely that chameleons are drawn to vineyards or at least perceive them as a suitable habitat. However, even in the absence of an inherent preference for vineyards, a third reason exists to believe chameleons are actively drawn to vineyards: given the mortality experienced following the harvest (see below), chameleons are likely drawn to vineyards repeatedly each season, due to differences in the density of chameleons inside versus outside vineyards. It could be hypothesized that chameleons are not drawn to vineyards but are pushed into them due to competitive exclusion. In such a case, it is expected that chameleons in vineyards would exhibit poorer body condition and that females would contain smaller clutches. In comparison to clutch sizes reported for this species from Spain (Díaz-Paniagua *et al*. 2002), the clutch size we found in female chameleons collected from the waste pile in the winery was larger (13.83 versus 17.43, Fig. 2, p = 0.02). Regarding body condition, according to data of *C. chameleon musae* in Israel (Sagi & Bouskila 2025), from approximately July onwards, males lose weight due to their continued investment in courting of receptive females (Cuadrado 2001). Females, in contrast, initially gain weight as they continue to feed but subsequently lose some weight (see Appendix 1). Unfortunately, we were unable to obtain data regarding body condition for *C. chamaeleon recticrista* from natural habitats during September. Hence, a direct comparison between the body condition of the chameleons we collected at the winery to the body condition of chameleons from natural habitats was not possible. However, our comparison between the body condition of chameleons surveyed in July (Bar-Yaacov *et al*. 2015) to the body condition of the chameleons we collected from the winery in September is consistent with the trend observed in *C. chamaeleon musae* (see Fig. 3).

**Figure 2.**
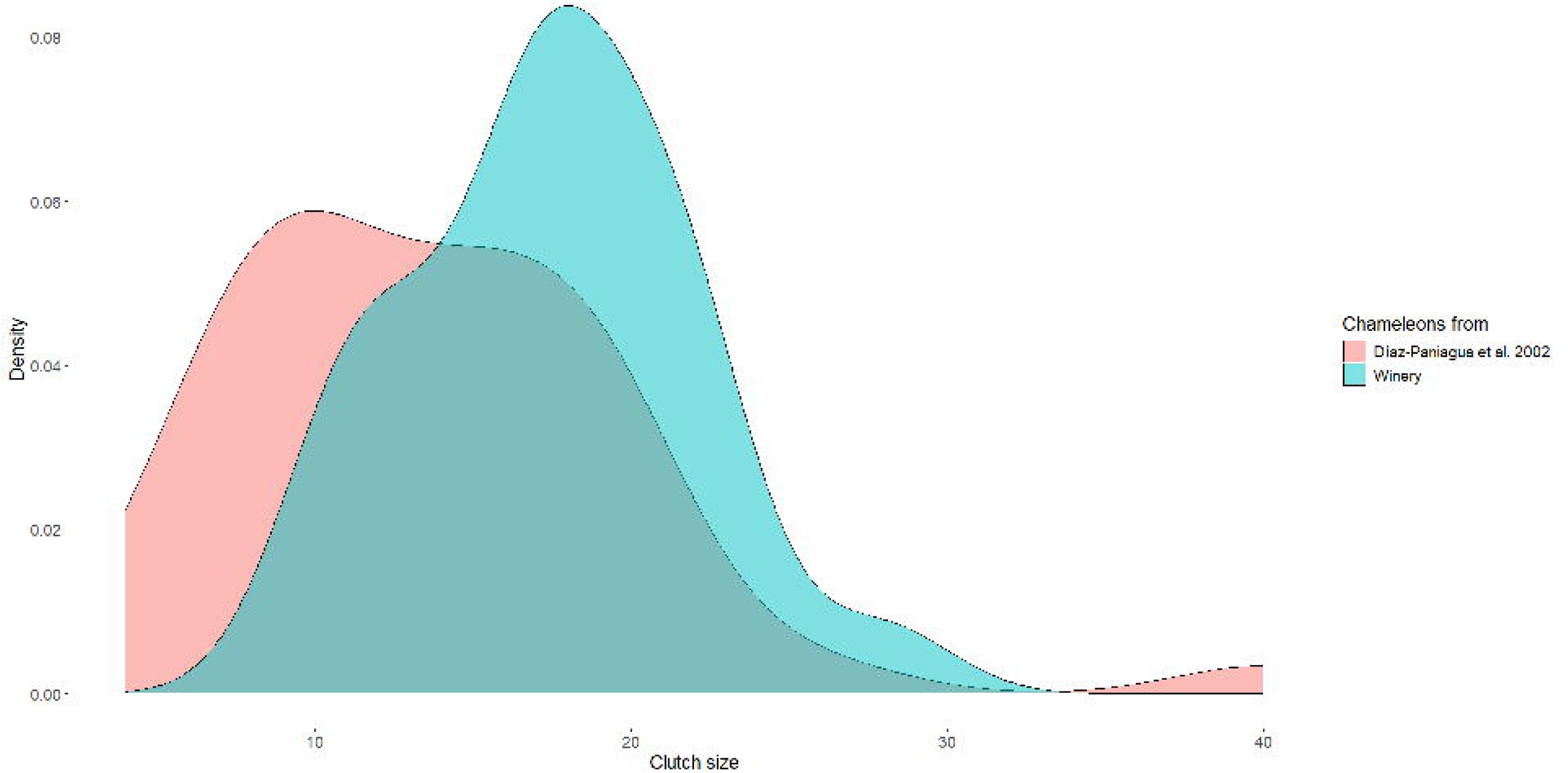
Density plot comparing clutch size from Diaz-Paniagua et al. (2002) versus the chameleons collected from the winery, using Kernel density estimate.

**Figure 3.**
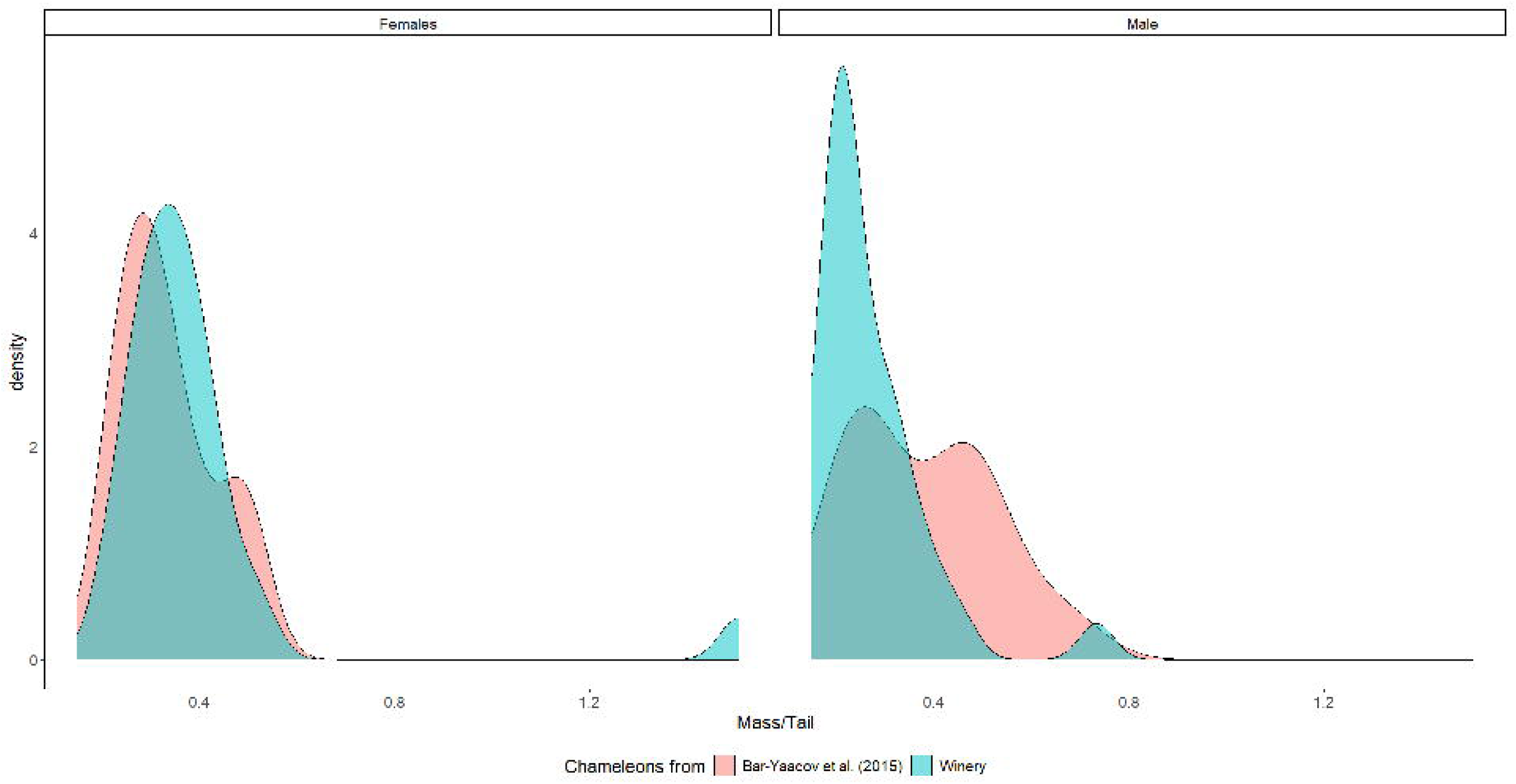
Density plot of body condition (mass/tail) comparing chameleons from Bar-Yaacov et al. (2015) versus chameleons collected at the winery, using Kernel density estimate.

Importantly, all but one of the female chameleons we collected at the winery (96%), contained unripe eggs. It has been suggested (Werner 2016) that chameleons belonging to this species typically reproduce only once during their lifetime, hence the entire reproductive potential of these chameleons (a total of 404 eggs) was most likely lost due to their accidental harvest in the vineyard.

To assess the possible effects of this hypothesized ecological trap, it is vital to consider the phenomenon we described at the population level. Since the chameleons in our data represent only the individuals harvested, we cannot directly determine their percentage of the entire population in the vineyard. However, there are several factors that should be considered in this context. First, chameleons rely primarily on camouflage as a means for avoiding danger rather than actively fleeing, and therefore, seem unlikely to escape mechanical harvest. Second, harvest is typically conducted during the night, when chameleons are inactive and sleep on branches. Third, from our personal experience in the field, chameleons at night tend to react slowly to noise and light. Furthermore, even when they do respond to a disturbance, their fleeing method often comprises dropping to the ground (Cuadrado *et al*. 2001, Pers. Obs.), which in the case of mechanical harvest would result in being collected by the harvest machine. Hence, we believe it is quite likely that most, if not all, chameleons in mechanically harvested vineyards are harvested. We found substantial variation in density between vineyards, even between some in close proximity (1.25 versus 6.36 chameleons/ha within < 10 km). This variation may be explained by several non-mutually exclusive possibilities. First, in certain cases, this variation may represent differences in the densities of surrounding populations. Second, differences in management practices such as the use of pesticides, tillage, etc., may affect the density of chameleons in a vineyard. Third, if chameleons are attracted to the grapes themselves, it is possible that the grape variety affects the vineyards’ attractiveness.

We were unable to obtain population density data of chameleons from the vineyards’ regions in natural habitats. The only available data were of chameleons from Samos, Greece (Dimaki *et al*. 2015). This comparison revealed that our estimated average density is lower than that reported from Samos (1.9 ind/ha versus 5.26 ind/ha respectively). However, as aforementioned, our results contain substantial variation such that some plots were closer to the estimation from Samos (one plot contained higher density) and may be influenced by the factors mentioned above.

The density of reptiles in vineyards may also be influenced by edge effects (reviewed by Ries *et al*. 2017). Small vineyards have larger circumferences relative to their area in comparison to larger vineyards, meaning, for example, that the proportion of vineyard potentially drawing animals from adjacent natural patches is larger. Biaggini & Corti (2021) reported that *Podarcis siculus* lizards were more abundant at vineyard borders, which may be the case for other species of reptiles as well. Therefore, it could be expected that small vineyards, such as those common in much of Europe (see section 1: introduction) would constitute more severe ecological traps. However, it should also be noted that smaller vineyards are more likely to be harvested manually (Domingues & Aguila 2016), in which case, the fitness costs for reptiles inhabiting the vineyards may be substantially lower.

Previously, ecological traps have been categorized according to a preference axis, such that equal preference traps are distinguished from ‘severe traps’, in which the ecological trap is *preferred* over the natural habitat (Hale & Swearer 2016). The reason that severe traps are regarded as such is that in the case of a preference for the low-quality habitat, the impact on the population is expected to be much higher due to strong source-sink dynamics. We suggest that mortality may constitute an additional useful axis along which ecological traps may be considered. Even in the case of an equal preference trap, when the mortality is very high, the effect on the population may be more severe than a ‘severe trap’ in which the fitness difference between the trap and natural habitat is lower. Furthermore, in the case of extreme mortality, the source-sink dynamics are expected to be particularly strong, even in the absence of an inherent preference for the trap habitat. We suggest that ‘severe preference traps’ should be distinguished from ‘severe fitness traps’. For the chameleons we described, it is likely that mechanically harvested vineyards constitute a ‘severe fitness trap’, regardless of whether they are inherently attracted to vineyards.

The interplay between specific agricultural practices and the focal species’ life history may affect the severity of an ecological trap substantially. Therefore, it is interesting to consider such differences between chameleons and the second most abundant species we collected from the winery (accounting together for >95% of reptiles collected). *Laudakia vulgaris* represents 27% of the reptiles found in the winery’s waste tank, with an average density of 0.83 ind/ha. Thus, vineyards seem to constitute an ecological trap for *L. vulgaris*, which may be attracted to vineyards for similar reasons as chameleons. However, there are several reasons to hypothesize that agamid mortality may be lower than chameleon mortality (relative to population size). First, while agamids, like chameleons, are inactive at night, chameleons tend to sleep at the ends of branches, while agamids tend to sleep on thick branches closer to the main stem (Pers. Obs.). Therefore, agamids may be less affected by the vine-shaking during the mechanical harvest process. Second, while chameleons seem to sleep almost exclusively on branches, agamids are known to find refuge in cracks and crevices found in rocks and trees, beneath rocks, and so on. Furthermore, when sheltering agamids are threatened, they inflate themselves and rely on their rough scales to render their extraction from the shelter more difficult (Werner 2016). Hence, depending on the specific layout and structure of the vineyard, it is possible that some agamids may occupy shelters that are less affected by the harvest process. Third, female agamids in Israel typically lay eggs between April-July while hatchlings appear during July-October (Werner 2016). Hence, though chameleons and agamids are both lizards of similar size that inhabit vineyards, differences in their life histories and the ways in which they interact with specific agricultural practices may alter how each species is affected. While some agamids are certainly killed during the process of mechanical harvest, the population-level effects may be less severe in their case. With the two axes of ecological traps in mind, vineyards may constitute both a ‘severe preference trap’ and a ‘severe fitness trap’ for chameleons, but only a ‘severe preference trap’ for agamids.

In addition to the chameleons and agamids discussed above, several other species are represented in our data. However, their small sample sizes hamper our ability to formulate any meaningful population-level generalizations regarding them. Notwithstanding, we point out that while adults of *C. chamaeleon recticrista* and *L. vulgaris* are of similar body size, *Mediodactylus orientalis* geckos and *Phoenicolacerta Laevis* lizards are substantially smaller and may therefore have been detected less during our surveys. The *Natrix tessellata* collected was a hatchling and therefore also smaller than the chameleons and agamids, but in any case, this species is not considered to be arboreal, and we do not consider it likely that it is substantially affected by the harvest process in vineyards (though other snake species may be affected to a greater extent). We also collected three Savigny’s treefrogs from the winery’s waste tank. Known to be arboreal and rely on lush vegetation, their presence in vineyards is unsurprising, particularly when the surrounding environment is dry and hot. However, also in this case, it is quite possible that their small size hampered the detection of additional individuals.

The phenomenon we described is unlikely to be limited to chameleons and agamids in vineyards. Many permanent crops may present similar problems for chameleons and agamids as well as for other species elsewhere. In general, any permanent crop harvested mechanically by shaking the entire tree and collecting the dislodged fruit immediately would likely result in high mortality of arboreal species inhabiting the trees. The severity of such agricultural practices will be determined by the interplay between the specific species and practices at hand: whether or not the inhabiting species are likely to escape the harvest process (i.e., mortality rate); whether or not the species at hand are actively drawn to the crop or sporadically inhabit it (i.e., preference, though as aforementioned, even in the absence of inherent preference source-sink dynamics may be formed); the timing of harvest relative to important life stages such as reproduction; whether or not collection of dislodged fruit occurs immediately after shaking or is delayed, potentially enabling fallen animals to escape, etc.

Importantly, the ecological effects of any given crop type might be substantially altered depending on the exact cultivation methods, harvest, etc. For example, in the case of vineyards, an alternative method employed primarily in small vineyards is to harvest the grapes by hand. This is more costly and labor-intensive but preferred for high-quality wines as it enables selectivity when picking the grapes. Compared to mechanical harvest, this method is unlikely to cause high mortality of arboreal reptiles, if any. Identifying species at risk could facilitate tailoring species-specific solutions based on the focal species’ life history. For example, it may be possible to ‘warn’ certain species (using noise, vibrations, light, scents, etc.) prior to commencement of the harvest.

In conclusion, alongside the negative effects of agriculture on biodiversity, various approaches can mitigate some of these effects and even confer certain positive effects on the environment. Such practices may prove invaluable for farmers, too, as they may increase benefits received through ecosystem services such as pollination, pest control, etc. However, even ‘sustainable’ practices may cause detrimental effects, which may differ greatly between crops, practices, and species. Specifically, we emphasize the importance of considering the various ways in which agricultural practices may unintentionally produce ecological traps, potentially risking not only biodiversity within the agricultural landscape but also depleting populations of species found in surrounding natural habitats. Furthermore, we suggest that distinguishing between ‘severe preference traps’ and ‘severe fitness traps’ may prove useful for mitigating the effects of ecological traps. Depending on the nature and mechanism of the trap, it may be possible to reduce its attractiveness (i.e., the preference) or adjust the specific methods to reduce the mortality it causes (i.e., the fitness costs).

All authors declare that they have no conflicts of interest.

## Acknowledgements

We thank Oded Berger-Tal, Liron Shani and Udi Gliksman and the Israel Nature and Parks Authority (INPA) for their insightful comments and their help in facilitating this study. We are indebted to the winery staff and vine growers who welcomed us into their facilities and enabled this study to take place.

This study was supported by grants from the Israel Science Foundation (ISF 1826/20), the U.S.-Israel Binational Science Foundation (BSF 1826/20), and the Minerva Center for the Study of Population Fragmentation. L.S. is funded by the Goldman Sonnenfeldt School of Sustainability and Climate Change, Ben Gurion University of the Negev.

During the preparation of this work the authors used ChatGPT in order to improve language and readability. After using this tool, the author reviewed and edited the content as needed and take full responsibility for the content of the publication

## Ethics statement

This study was conducted under the approval of the Israel Nature and Parks Authority (permit number 2021/42820).

