## Supplementary material for "The double-edged sword of wildlife biodiversity in the agricultural matrix: a case study of reptiles in vineyards": Online Appendices

Appendix 1:

Based on data of *Chamaeleo chamaeleon musae* from the Negev desert (Authors, accepted), we examined the trends in body size of males and females, starting at the beginning of the mating season and through the reproductive season. These data were measured between 2009-2021 and represent 507 observations relevant to our comparison between months.

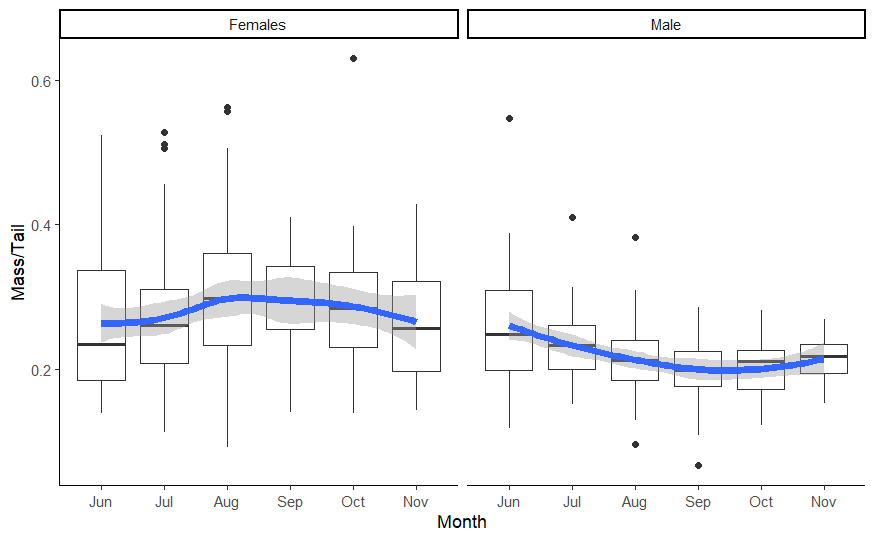

Figure A1. Change in body condition throughout the reproductive season for males and females separately, the blue line represents local regression (LOESS, span = 0.75)

References

Authors. (accepted). First evidence of yearly allochrony in a terrestrial vertebrate: A case study of an annual chameleon. *Ecology*.

Appendix 2:

Table A1: Density of chameleons and agamids from vineyards in different areas.

The ‘Mediterranean – Total’ includes all chameleons and agamids, excluding those from unknown locations.
When possible, we separated the batches into grape varieties, using the following abbreviations: PV – Petit Verdot; CF - Cabernet Franc; CS - Cabernet Sauvignon; MER – Merlot.
* Judaean ‘Mountains – Total’ – includes one chameleon for which we could not distinguish between PV and CF batches.
† In this case, we could not distinguish between Shephelah and Galilee for chameleons, but were able to for Agamids.
‡ No chameleons were found in batches from Ramon, which was excluded from the total calculation since Ramon is outside of the chameleons’ distribution.

| date | Vineyards’ location | Vinyards’ area (ha) | Chameleons | Agamids | Chameleons / ha | Agamids / ha |
| --- | --- | --- | --- | --- | --- | --- |
| 17/08/2021 | Unknown |  | 5 |  |  |  |
| 01/09/2021 | Shephelah - PV | 3.1 | 3 | 0 | 0.967741935 |  |
| 01/09/2021 | Shephelah | 4.7 | † | 1 |  | 0.212766 |
| 01/09/2021 | Galilee - CS | 2 | † | 14 |  | 7 |
| 01/09/2021 | Shephelah and Galilee | 6.7 | 17 | † | 2.537313433 |  |
| 01/09/2021 | Ramon | 4 | ‡ | 1 |  | 0.25 |
| 19/09/2021 | Northern highlands (Galilee and Golan) | 5.24 | 9 | 2 | 1.717557252 | 0.381679 |
| 10/09/2023 | Unknown |  | 5 |  |  |  |
| 10/09/2023 | Mediterranean - CS | 2.4 | 1 | 0 | 0.416666667 | 0 |
| 12/09/2023 | Judaean Mountains - CF | 1.1 | 7 | 4 | 6.363636364 | 3.636364 |
| 12/09/2023 | Judaean Mountains - PV | 2.4 | 3 | 0 | 1.25 | 0 |
| 12/09/2023 | Judaean Mountains - Total | 3.5 | 11* | 4 | 3.142857143 | 1.142857 |
| 12/09/2023 | Galilee - MER | 1.3 | 4 | 0 | 3.076923077 | 0 |
| 12/09/2023 | Shephelah | 1.5 | 5 | 0 | 3.333333333 | 0 |
| 12/09/2023 | Mediterranean | 6.1 | 14 | 5 | 2.295081967 | 0.819672 |
| 12/09/2023 | Unknown |  | 3 | 1 |  |  |
|  | **Mediterranean - Total** | **33.84** | **64** | **28** | **1.891252955** | **0.827423** |
